# Photocatalytic plant LPOR forms helical lattices that shape membranes for chlorophyll synthesis

**DOI:** 10.1101/2020.08.19.257774

**Authors:** Henry C. Nguyen, Arthur A. Melo, Jerzy Kruk, Adam Frost, Michal Gabruk

## Abstract

Chlorophyll (Chl) biosynthesis, crucial to life on Earth, is tightly regulated because its precursors are phototoxic^1^. In flowering plants, the enzyme Light-dependent Protochlorophyllide OxidoReductase (LPOR) captures photons to catalyze the penultimate reaction: the reduction of a double-bond within protochlorophyllide (Pchlide) to generate chlorophyllide (Chlide)^2,3^. In darkness, LPOR oligomerizes to facilitate photon energy transfer and catalysis^4,5^. However, the complete 3D structure of LPOR, the higher-order architecture of LPOR oligomers, and the implications of these self-assembled states for catalysis, including how LPOR positions Pchlide and the cofactor NADPH, remain unknown. Here we report the atomic structure of LPOR assemblies by electron cryo-microscopy (cryoEM). LPOR polymerizes with its substrates into helical filaments around constricted lipid bilayer tubes. Portions of LPOR and Pchlide insert into the outer membrane leaflet, targeting the product, Chlide, to the membrane for the final reaction site of chlorophyll biosynthesis. In addition to its crucial photocatalytic role, we show that in darkness LPOR filaments directly shape membranes into high-curvature tubules with the spectral properties of the prolammelar body, whose light-triggered disassembly provides lipids for thylakoid assembly. Our structure of the catalytic site, moreover, challenges previously proposed reaction mechanisms^6^. Together, our results reveal a new and unexpected synergy between photosynthetic membrane biogenesis and chlorophyll synthesis in plants orchestrated by LPOR.

## Introduction

Chlorophyll (Chl) is the major light-harvesting molecule on our planet. Its biosynthesis involves the generation of progressively more efficient light-absorbing, yet also phototoxic, pigments with increasingly hydrophobic properties^1^. The complete synthetic pathway requires an efficient flow of intermediates between biosynthetic enzymes, coordination between pigment production and photosynthetic membrane formation, and a regulatory mechanism to synchronize the chloroplast-based synthesis of the pigment with nuclear expression of photosynthetic proteins.

In oxygenic phototrophs, Chl production depends on photons because the Light-dependent Protochlorophyllide OxidoReductase (LPOR, E.C. 1.3.1.33) is a photoactivated reductase. This enzyme produces Chlide^3^, the final tetrapyrrole ring that harnesses the lower-energy wavelengths of sunlight more efficiently than its precursors^7^. LPOR is one of only a few known light-activated enzymes and its mechanism of action remains elusive^6^. In the darkness, plant LPOR accumulates in immature chloroplasts, or etioplasts, within a paracrystalline cubic lattice of branched tubules called prolamellar bodies (PLBs)^3^. PLBs comprise the same lipids as photosynthetic membranes, including predominantly monogalactosyldiacylglycerol (MGDG), digalactosyldiacylglycerol (DGDG), phosphatidylglycerol (PG) and sulfoquinovosyldiacylglycerol (SQDG)^8^. Upon illumination, LPOR catalyzes the reduction of Pchlide within milliseconds as PLBs simultaneously disassemble. This coupled reductase activity and membrane structure dismantling is the first step in the formation of thylakoid membranes^9^.

Decades of investigation have revealed that LPOR forms oligomers within PLBs, enabling efficient light energy transfer between Pchlides within the complex (summarized in^4,8^). Recent in vitro work showed that PG and SQDG increase the NADPH binding affinity to plant LPOR, while MGDG affects the spectral properties of the complex and may trigger oligomerization^5^. To better understand these effects, we investigated the structural interactions between the enzyme, Pchlide, NADPH, and the specific PLB lipids that support oligomerization and enhance catalytic activity.

### Plant LPORs form filaments with lipids and substrates

We report that all three *A. thaliana* LPOR isoforms (AtPORA, AtPORB, AtPORC) form helical tubes in darkness in the necessary presence of Pchlide, NADPH, and liposomes containing at least MGDG and PG (Fig. S1a). These assemblies can be microns long with diameters that range from ∼10 nm to ∼30 nm, depending on the LPOR isoform, the ratios of the lipid species included in the liposomes, and the relative lipid:enzyme concentrations. The oligomerization process is fast, with an upper bound of 5 minutes at room temperature, and requires darkness.

AtPORB produced the most regular tubes. We, therefore, optimized the liposome composition and reaction conditions for this isoform to generate samples for high-resolution imaging. While the optimized lipid composition (50 mol% MGDG / 35 mol% DGDG / 15 mol% PG) differs from the relative composition of the PLB found in plants, these AtPORB filaments display similar morphologies to those produced with liposomes closely mimicking chloroplast inner membranes^10^ and PLBs^11^ (Fig. S1b). Moreover, tubes produced with either the optimized lipid mixture or chloroplast inner membrane-like and PLB-like liposomes generated identical fluorescence emission spectra for Pchlide. Importantly, the emission maxima in all cases were identical to those characteristic of isolated PLBs^12^ (Fig. S1c).

**Fig. S1.**
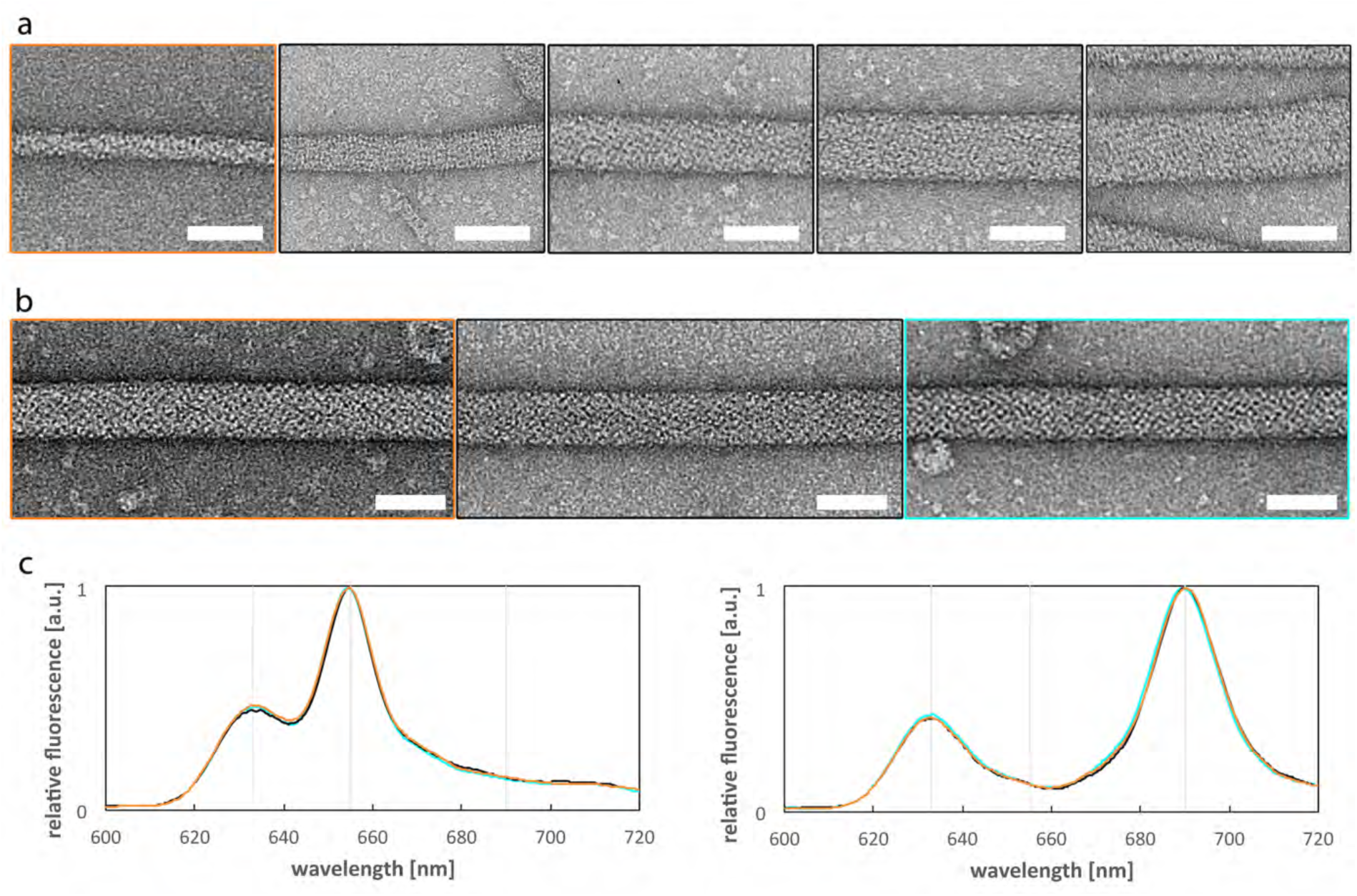
Plant LPOR can form filaments of different sizes. **a**, Negative stain electron microscopy (EM) micrographs of AtPORB filaments with different diameters formed from liposomes composed of 2.5 mol% MGDG / 50 mol% DGDG / 47.5 mol% PG (orange frame) or 50 mol% MGDG / 25 mol% DGDG / 25 mol% PG (black frames). Scale bar: 25 nm. **b**, Negative stain EM micrographs of AtPORB filaments produced from liposomes mimicking the lipid composition of PLBs (52 mol% MGDG / 32 mol% DGDG / 8 mol% PG, 8 mol% SQDG; orange frame, chloroplast inner membranes (50 mol% MGDG / 30 mol% DGDG / 8 mol% PG, 5 mol% SQDG, 7 mol% PC, cyan frame) and our optimized membrane composition (50 mol% MGDG / 35 mol% DGDG / 15 mol% PG, black frame). Scale bar: 25 nm. **c**, Low-temperature fluorescence emission spectra before (left panel) and after illumination (right panel) of the filaments produced with the liposomes mimicking the lipids of PLBs (orange), chloroplast inner membrane (cyan) and for our optimized membrane composition (black). Grey vertical lines indicate the emission maxima of: free Pchlide (632 nm), PLBs or the filaments (655 nm) and Chlide (690 nm).

### Pchlide is embedded within the AtPORB lattice and the outer leaflet of the membrane

We next analyzed the optimized AtPORB assemblies by electron cryo-microscopy (cryoEM). The tubes were flexible and heterogeneous in diameter, yet we succeeded in classifying a well-ordered subset that yielded a 3D reconstruction to ∼3.1 Å (FSC 0.143 half-map criterion, Fig. 1b,c, Table S1, Fig. S2, and Video S 1). This reconstruction revealed AtPORB antiparallel dimers assembled into a strand, with four such strands, or a ‘4-start’ assembly, encasing a membrane bilayer tubule (Fig. S3). The overall diameter of the filament was ∼24 nm, and the distance between outer leaflet headgroups was ∼12 nm versus ∼6 nm between the inner leaflet headgroups. Such a thin bilayer thickness, just ∼3 nm, is consistent with previous observations of highly constricted membranes^13^.

**Fig. 1.**
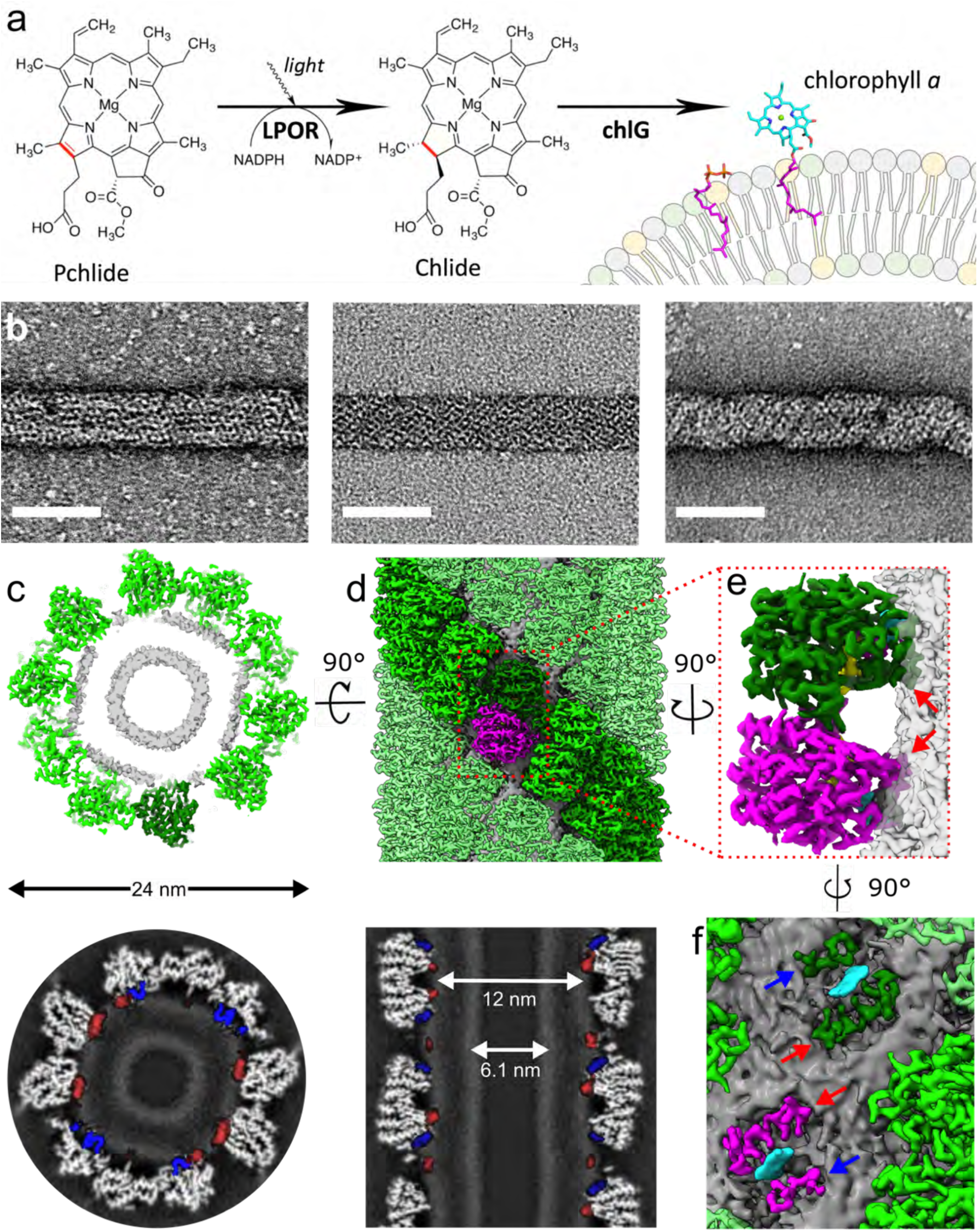
Plant LPORs form filaments around a membrane bilayer. **a**, The last steps of Chl biosynthesis in flowering plants. In the penultimate reaction, LPOR reduces a specific double bond of Pchlide upon illumination. In the final reaction, Chlide is then esterified by Chl synthase (chlG) with a phytol moiety to produce Chl. **b**, Negative stain electron microscopy (EM) micrographs of AtPORA (left), AtPORB (middle), and AtPORC (right) filaments. Scale bar: 25 nm. **c**, The cryoEM 3D reconstruction of the membrane-bound AtPORB filament. Top-down view in surface representation (top) and a grey-scale slice (bottom). **d**, Exterior view (top) and grey-scale slice (bottom) along the helical axis. AtPORB (green) coats the exterior of the membrane bilayer (grey). An AtPORB dimer is highlighted with each protomer in dark green or magenta. In the grey-scale slices, helix α10 and the Pchlide loop from all protomers are false-colored red and blue, respectively. Diameters of the entire tube and the membrane leaflet peak-to-peak distances are annotated. **e**, The AtPORB dimer partially inserts into the outer leaflet. NADPH and Pchlide are colored in yellow and cyan, respectively. Red arrows denote helix α10. **f**, Zoomed in view of the membrane-embedded Pchlide with helix α10 and the Pchlide loop labeled (blue arrows).

Overall, this reconstruction shows that plant LPOR is itself a direct membrane-binding and membrane-remodeling enzyme that assembles to position its substrate, Pchlide, within a helical lattice embedded in the outer leaflet of the membrane. Notably, two hydrophobic segments of each protomer insert into the outer leaflet: a region of the LPOR-specific loop we name “the Pchlide loop” (residues 229-243) and helix α10 (Fig. 1d). Between these sections, we observed apparent density for Pchlide, which was also partially embedded in the outer leaflet (Fig. 1e). In addition to coordinating substrate binding, the Pchlide loop and helix α10 likely help target AtPORB to the bilayer, increasing its local concentration to trigger polymerization, inducing positive curvature membrane remodeling, and, consequently, membrane tubulation.

**Fig. S2.**
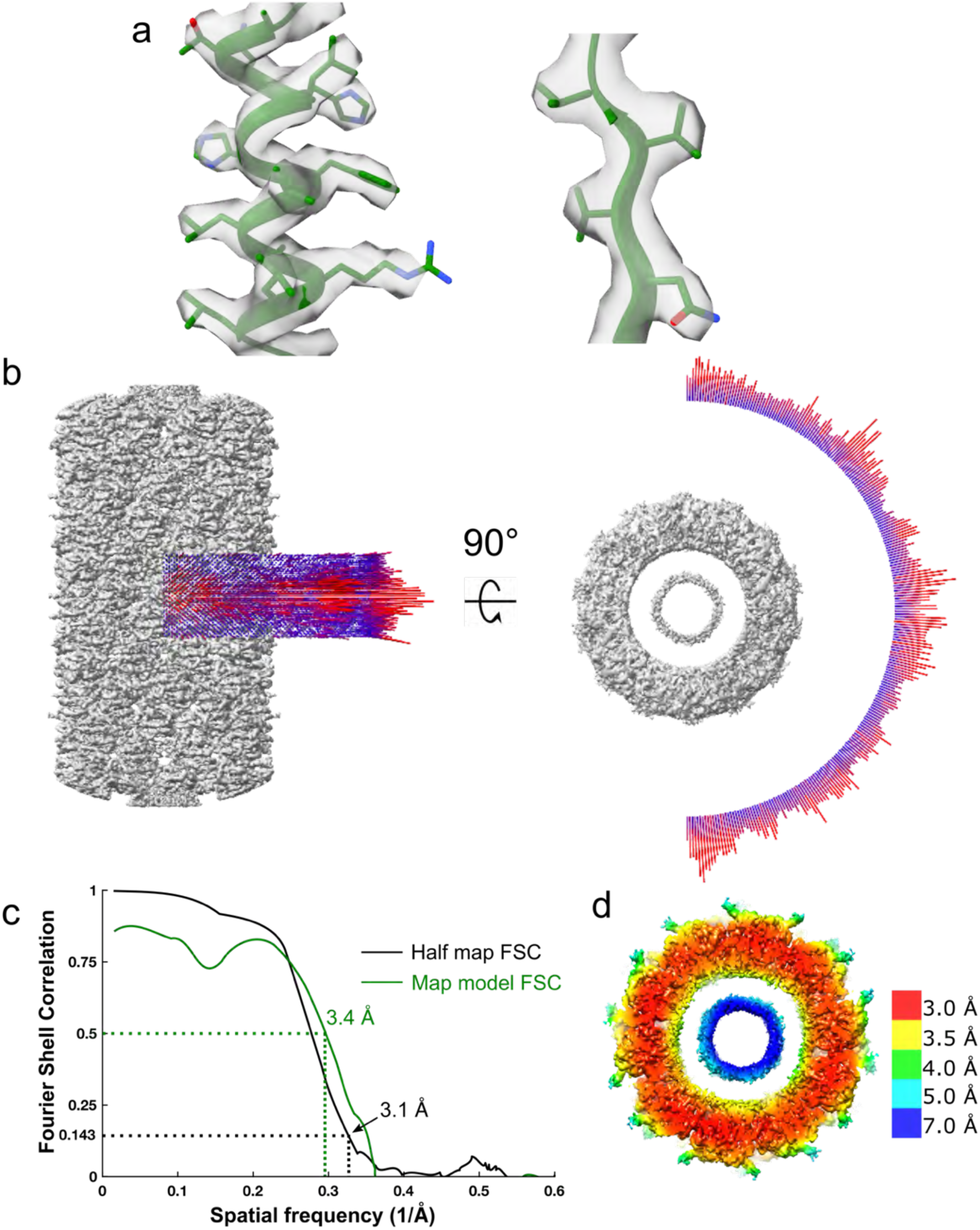
CryoEM validation of the AtPORB filament. **a**, Representative fit of the cryoEM density to the atomic model. **b**, Angular distribution of the membrane-bound AtPORB cryoEM 3D reconstruction. **c**, Independent half-map and map-model Fourier shell correlation (FSC) of the reconstruction. **d**, Local resolution estimates.

**Fig. S3.**
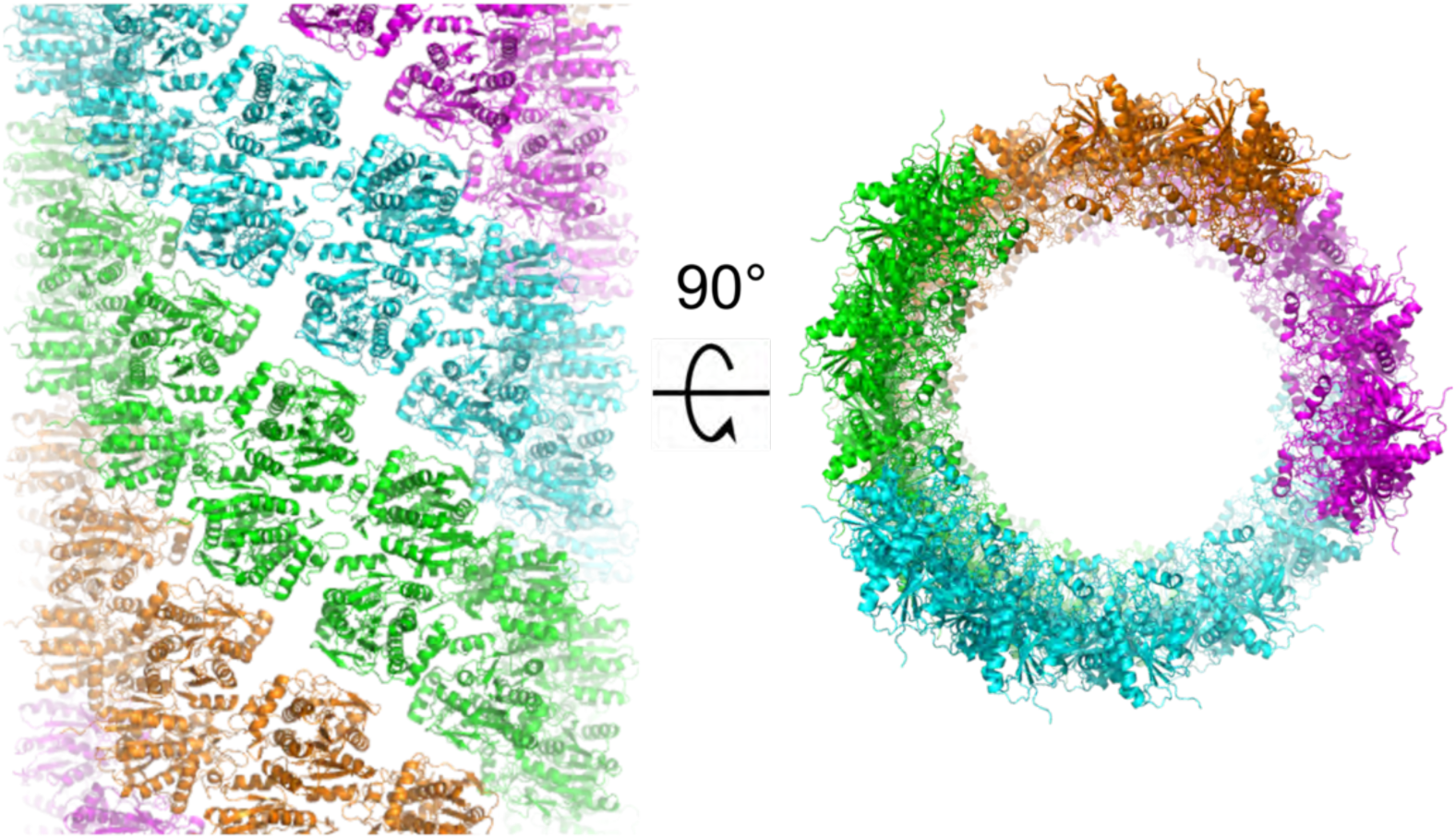
A 4-start helix of plant LPOR. Exterior and top-down views of the 4-start PORB helix. Each start comprises a strand of antiparallel dimers and colors in orange, green, cyan, or magenta.

**Video S1.**
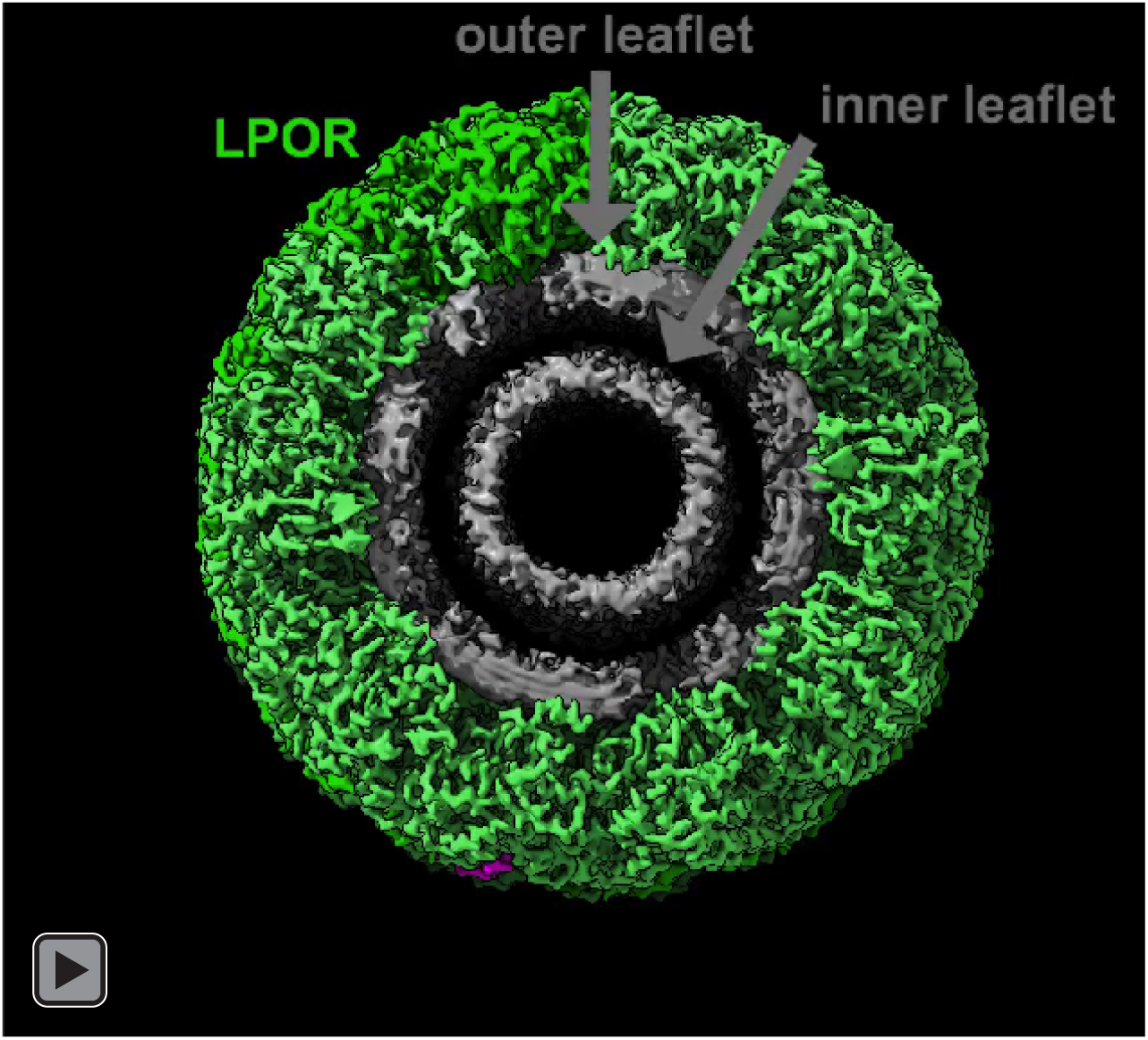
Structure of the membrane-bound LPOR filament with substrates bound.

**Table S1.** CryoEM data collection, refinement, and validation statistics.

|  |  |
| --- | --- |
|  | Membrane-bound LPOR<br>(EMD-22364,<br>PDB 7JK9) |
| **Data collection and processing** |  |
| Magnification | 105,000 |
| Voltage (kV) | 300 |
| Electron exposure (e<sup>-</sup>/Å<sup>2</sup>) | 73.5 |
| Defocus range (μm) | -0.2 to -1.9 |
| Pixel size (Å) |  |
| Collection | 0.835 |
| Processing | 1.11333 |
| Refined helical symmetry | 50.34°, 43.1 Å |
| Point group symmetry | C2 |
| Initial particle images (no.) | 294,156 |
| Final particle images (no.) | 17,898 |
| Map resolution (Å) | 3.1 |
| FSC threshold | 0.143 |
| Map resolution range (Å) | 10-2.9 |
| **Refinement** |  |
| Initial model used (PDB code) | De novo |
| Model resolution (Å) | 3.4 |
| FSC threshold | 0.5 |
| Map sharpening *B* factor (Å<sup>2</sup>) | -97 |
| Model composition |  |
| Nonhydrogen atoms | 101,480 |
| Protein residues | 12,640 |
| Ligands | NADPH: 40<br>Pchl<sub>a</sub>: 40<br>MGDG: 40 |
| *B* factors (Å<sup>2</sup>) |  |
| Protein | 44 |
| Ligands | 50 |
| R.m.s. deviations |  |
| Bond lengths (Å) | 0.007 |
| Bond angles (°) | 1.289 |
| **Validation** |  |
| MolProbity score | 1.26 |
| Clashscore | 4.99 |
| Rotamer outliers (%) | 0.39 |
| Ramachandran plot |  |
| Favored (%) | 98.09 |
| Allowed (%) | 1.91 |
| Disallowed (%) | 0 |

### Oligomerization of AtPORB arranges Pchlide for efficient energy transfer

Filament assembly is governed by interactions between three unique interfaces of AtPORB: the antiparallel dimer, the intra-strand, and the inter-strand interfaces (Fig. 2). Antiparallel dimers are the core building blocks of the filament, where the buried surface area (BSA) between two protomers (A and B) is ∼840 Å^2^. Hydrogen bonds (A:R128-B:Y186^backbone^, A:Q155-B:V142^backbone^) and cation-pi stacking interactions (F123-H144) further stabilize it (Fig. 2b). The antiparallel arrangement leads to a two-fold symmetry relating the protomers, so the symmetric interactions (B:R128-A:Y186) also occur. While Pchlide-free cyanobacterial LPORs crystallized as dimers in the asymmetric unit^6,14^, the interface is different from our antiparallel dimer and may be due to crystallization artifacts. Since there are only minor sequence differences between plant and cyanobacterial LPORs at this interface, it is possible that cyanobacterial LPORs dimerize in the same fashion as plant LPOR.

**Fig. 2.**
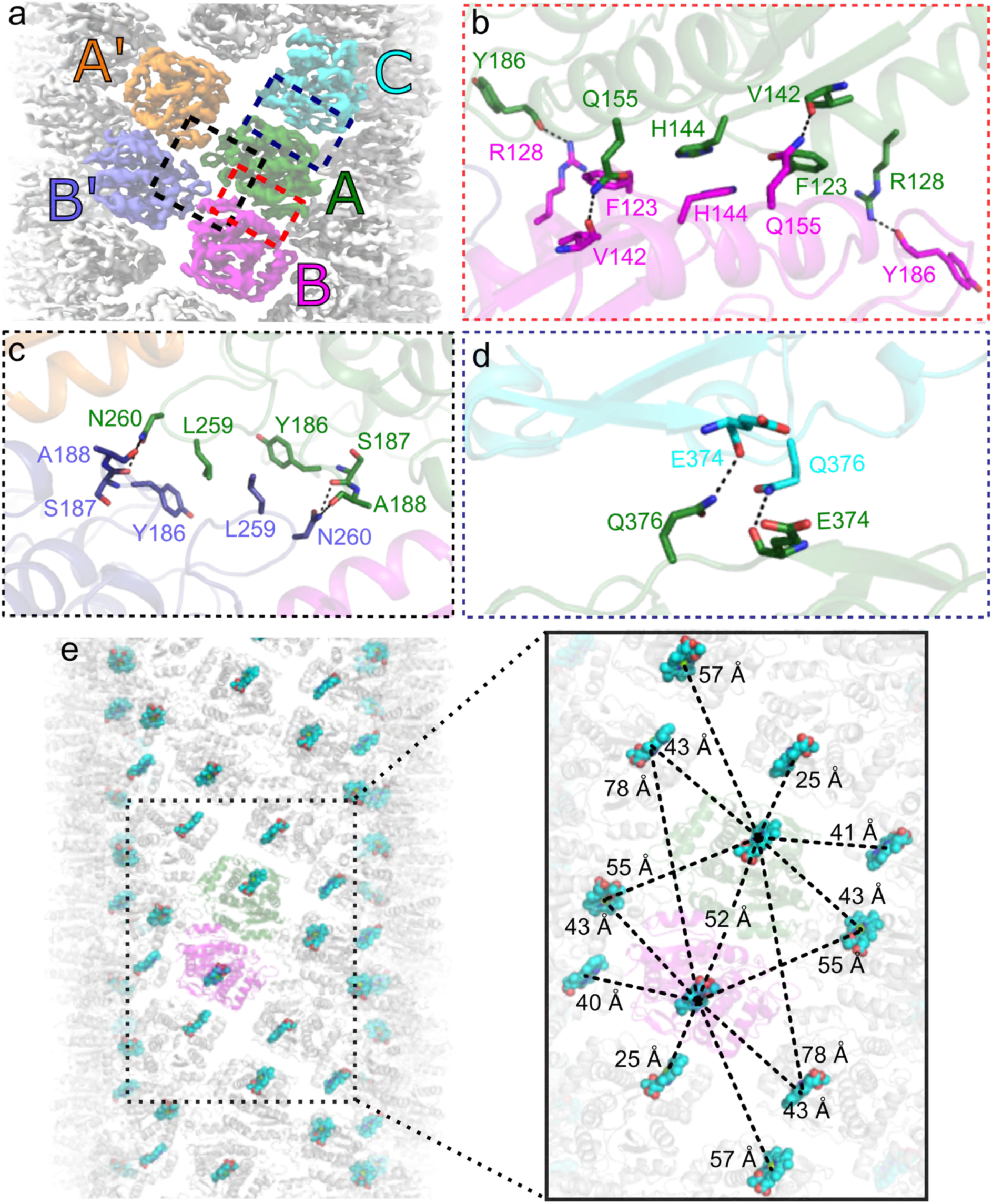
AtPORB oligomerization creates a periodic array of Pchlides. **a**, Exterior view of the AtPORB 3D reconstruction with protomers labeled. **b**, Protomers A and B constitute the antiparallel dimer interface, the red boxed area in **a. c**, The intra-strand interface formed from two antiparallel dimers, the black boxed area in **a**, with protomers A’ and B’. **d**, The inter-strand interface, the blue boxed area in **a. e**, Left, Exterior view of the filament as ribbons showing the arrangement of Pchlides (cyan spheres) within the filament, an AtPORB dimer is colored as in **a** while other subunits are in white. Right, close up view showing the distance between adjacent Pchlides.

Two LPOR dimers come together to form the intra-strand interface, and this mediates polymerization of the strands (a dimer of A-B with a dimer of A’-B’, Fig. 2c). Between A and A’, or the symmetry related B and B’, there is ∼370 Å^2^ of BSA. Between A and B’, there is ∼295 Å^2^ of BSA with two hydrogen bonds (A:N260-B’:S187^backbone^ and A:N260-B’:A188^backbone^). Due to the symmetry, the total BSA across all subunits is ∼1330 Å^2^, making this the largest interface driving polymerization. At the four-way junction where all four protomers meet, hydrophobic interactions occur across A:L259 and A:Y186 with B’:Y186 and B’:L259. Notably, the loop contributing to this junction involves a unique insertion compared to cyanobacterial LPORs^15^, allowing for increased BSA to promote oligomerization for plant LPORs (Fig. S4b). Recently, cyanobacterial LPORs have also been shown to form oligomers^6^. By negative stain EM, we see morphologically different cyanobacterial LPOR filaments in the presence of our optimized membrane composition, suggesting they may utilize a different polymerization interface (Fig. S4c).

The inter-strand interface, bringing strands together to form the full filament, is the weakest interface and driven by local two-fold symmetric interactions between protomers A and C. Only two hydrogen bonds (A:E374^backbone^-C:Q376 and the symmetry related interaction) and ∼280 Å^2^ of BSA are involved (Fig. 2d). However, when propagated across all the subunits within each strand, this stabilizes the whole tubule. Importantly, this interaction is likely malleable and this may explain the formation of different *n*-start filaments.

The sum of these interactions drive AtPORB oligomerization, leading to a helical array of Pchlides around the tubule (Fig. 2e). Adjacent Pchlides are between ∼25 to 78 Å away from each other and, importantly, these distances are well within the lengths necessary for efficient resonant energy transfer upon illumination. This notion is supported by the increased catalytic efficiency of in vitro^5^ oligomeric LPOR complexes compared to monomers. Notably, the shortest distance is from neighboring strands, suggesting that inter-strand formation is important and explains the higher reaction efficiency in its filamentous form^5^. Concurrently, it may also function to effectively reduce reactive oxygen species production upon illumination (a proposed function of the PLB summarized in ^4^), providing multiple acceptors of the absorbed energy and preventing diffusion of the phototoxic pigment into the stroma of chloroplasts.

**Figure S4.**
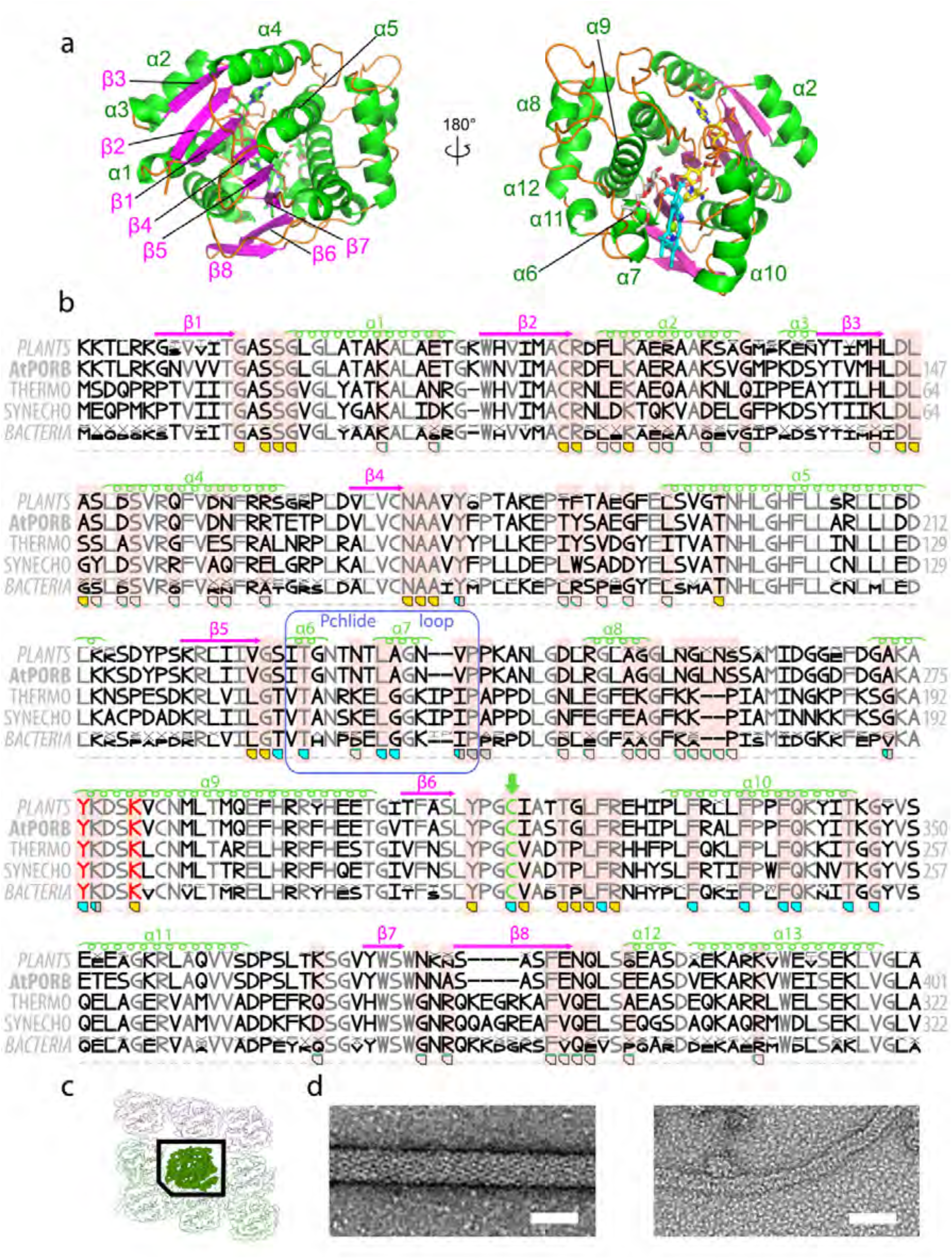
Comparison of plant and cyanobacterial LPORs. **a**, Secondary structure elements of AtPORB annotated onto the tertiary structure. **b**, The comparison of the sequences of AtPORB, *T. elongatu*s LPOR (Thermo), *Synechocystis* LPOR (Synech), and the consensus sequences of LPORs originating from plants and cyanobacteria (187 and 301 sequences, respectively). The size of the letter indicates the frequency of the given amino acid at the given position, while x indicates multiple amino acids with low frequency at a given position. Transit peptides of plant LPORs are excluded. Residues conserved in all sequences are grey. The secondary structure elements and Pchlide loop of AtPORB are marked. AtPORB residues known to play a role in oligomerization and substrate binding are highlighted with a pale red background. Orange/cyan/grey polygons indicate residues of NADPH/Pchlide/MGDG binding sites, respectively. Edges of the polygon highlighted in green indicate the oligomerization interfaces the given residue is involved in: bottom edge – antiparallel dimer, top edge – intra-strand interface, side edges – inter-strand interface. The previously proposed catalytic YxxxK motif is marked in red. Cysteine residue that we suggest to be a proton donor in the reaction is marked in green. **c**, Schematic representation of the LPOR interfaces for the polygons in **b. d**, Negative stain EM micrographs of AtPORB (left) and *Synechocystis* LPOR (right) filaments produced with our optimized membrane composition. Scale bar: 25 nm.

### LPOR-bound Pchlide is flanked by NADPH and MGDG

Within each protomer, AtPORB traps Pchlide between the Pchlide loop, helix α10, and the membrane (Fig. 1e). Pchlide binds by predominately hydrophobic interactions, with a few polar residues possibly responsible for the substrate specificity of LPOR, for example: Q331 is proximal to Mg^2+^ of the tetrapyrrole ring, H319 and Y177 are close to the carbonyl group of Pchlide, and Y276 is possibly interacting with the carboxyl group of the propionate moiety (Fig. 3a). In support of our Pchlide binding site, evolutionarily conserved residues that interact with Pchlide in our structure, namely AtPORB T230 and Y306, have been previously shown to decrease Pchlide binding when mutated in cyanobacterial LPORs^6,17,18^. While a binding site for Pchlide was proposed based on homology models and assumed proximity to NADPH^5,17^, our observation that Pchlide is partially embedded within the amphipathic region of the membrane’s outer leaflet is unexpected and novel.

**Fig. 3.**
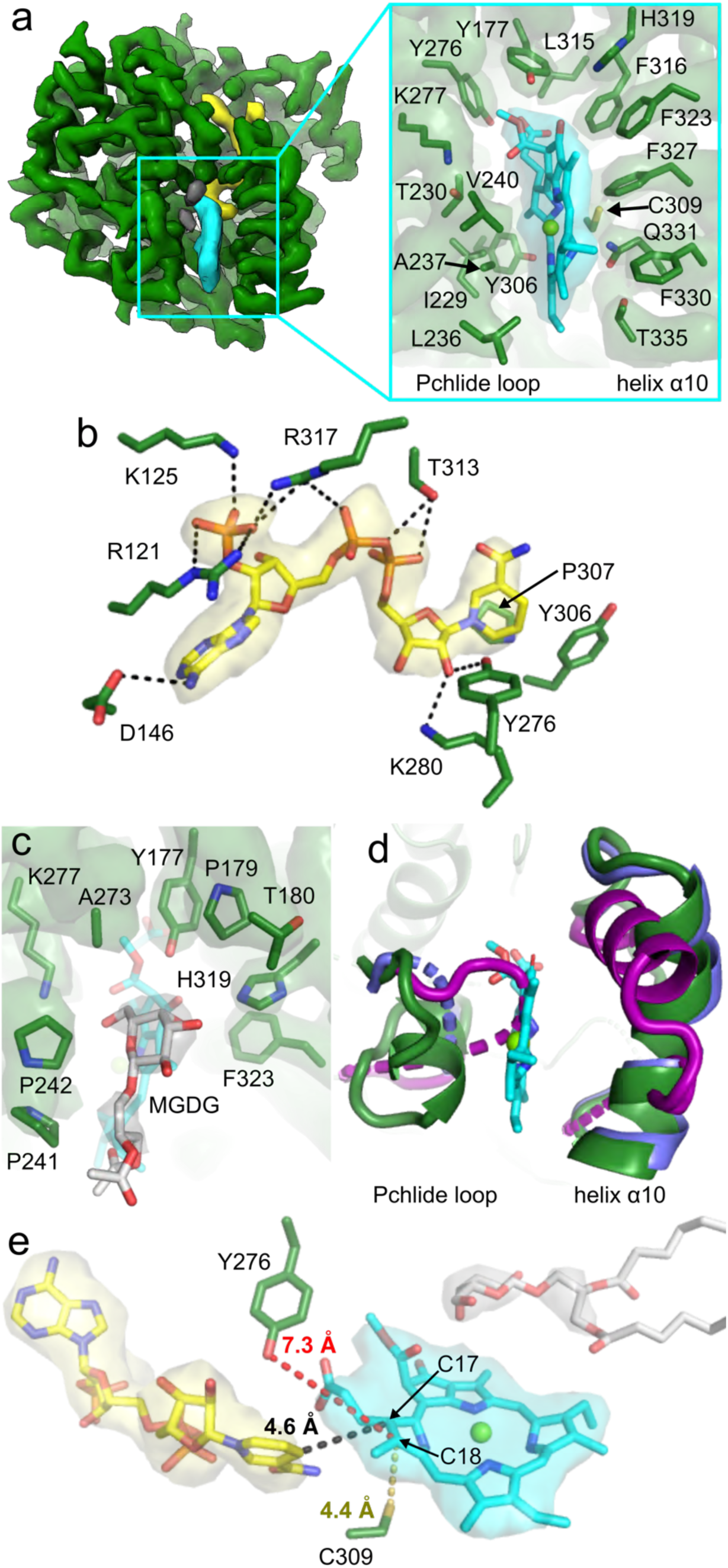
Detailed interactions between AtPORB and its substrates suggest a novel catalytic mechanism. **a**, Pchlide (cyan) is sandwiched between the Pchlide loop and helix α10. **b**, LPOR side chains (green sticks) involved in binding to NADPH (yellow). Dashed lines represent hydrogen bonds or salt bridges. **c**, The headgroup of MGDG (light grey) caps the edge of the Pchlide binding site. **d**, Binding of Pchlide induces structural changes. Superposition of AtPORB (dark green) to Pchlide-free LPOR structures from *T. elongatu*s (PDB: 6L1H, slate blue) and *Synechocystis* sp. (PDB: 6R48 or 6L1G, dark purple). **e**, Position of NADPH, Pchlide, and MGDG within AtPORB. Proposed catalytic residues are shown. CryoEM densities are represented as semi-transparent surfaces.

NADPH, clearly resolved, does bind adjacent to Pchlide in a mode similar to the cyanobacterial LPOR co-crystal structures with NADPH (Fig. 3b). While we could not resolve coordinated water molecules as in the crystal structures, there are some minor differences with respect to the NADPH interactions (Fig. S5). Previously, the lipid phosphatidyl glycerol (PG) was shown to dramatically increase the affinity of AtPORB towards NADPH^15^. We did not identify this lipid in our reconstruction despite its presence in the sample, possibly due to either its low occupancy or high mobility, but did find it was essential for plant LPOR to form filaments. Due to the acidic charge of the PG headgroup, it may help recruit the photoenzyme to the membrane through electrostatic interactions.

Surprisingly, we also observed an additional weak density between Pchlide and the Pchlide loop. The density is largely surrounded by hydrophobic residues and the keto group of Pchlide (Fig. 3c). Previously, infrared spectroscopy suggested that hydrogen bonds (with an unknown donor) to the keto group were responsible for the red-shift of the fluorescence emission maximum (FEM) of the pigment within LPOR^19^. We recently demonstrated that the lipid MGDG red-shifts the FEM^5^. Given that this density is located in the headgroup region of the membrane (Fig. 1e), we propose that it corresponds to the galactosyl headgroup, which fits well into the density (Fig. 3c). The lipid likely stabilizes pigment binding and plays an essential catalytic role because the red-shift of the Pchlide FEM decreases the energy of the transition between different excited states of the pigment^19^.

**Fig. S5.**
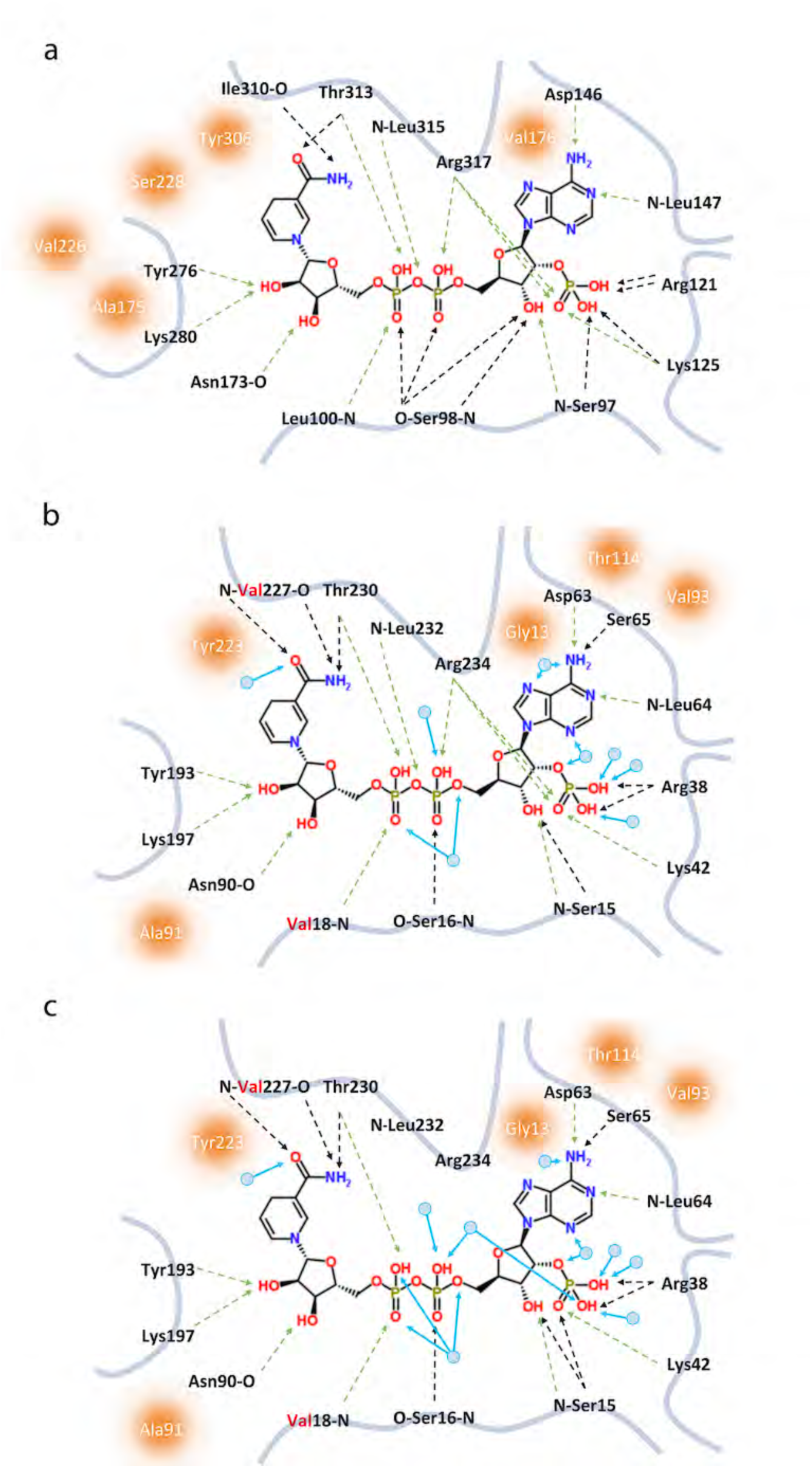
Comparison of the NADPH binding site across plants and cyanobacteria. **a**, NADPH binding pocket of AtPORB. **b**, NADPH binding pocket of LPOR form *T. elongatu*s (PDB: 6L1H). **c**, NADPH binding pocket of LPOR form *Synechocystis* (PDB: 6L1G). Hydrophobic interactions are shown as orange circles and hydrogen bonds are shown as dashed lines. Green dashed lines indicate the identical bonds between homologous residues. Resides in red indicate non-conserved residues. Waters are shown as small light blue circles.

**Fig. S6.**
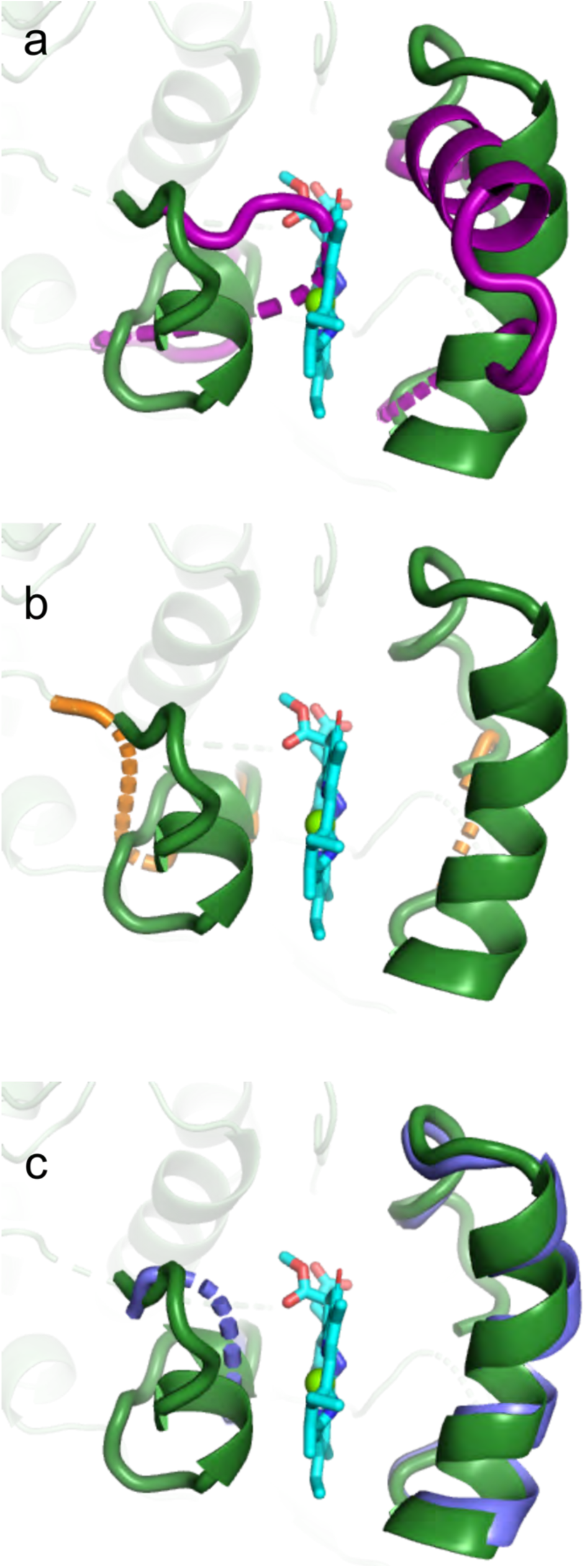
Structural comparison of Pchlide binding motifs between Pchlide-bound plant and Pchlide-free cyanobacterial LPOR. Superposition of Pchlide-bound AtPORB (dark green) with Pchlide-free LPOR from *Synechocystis* LPOR (**a**, dark purple, PDB: 6R48 or 6L1G), *T. elongatus* (**b**, PDB: 6RNW, orange), and *T. elongatu*s (**c**, PDB: 6L1H, slate blue).

**Video S2.**
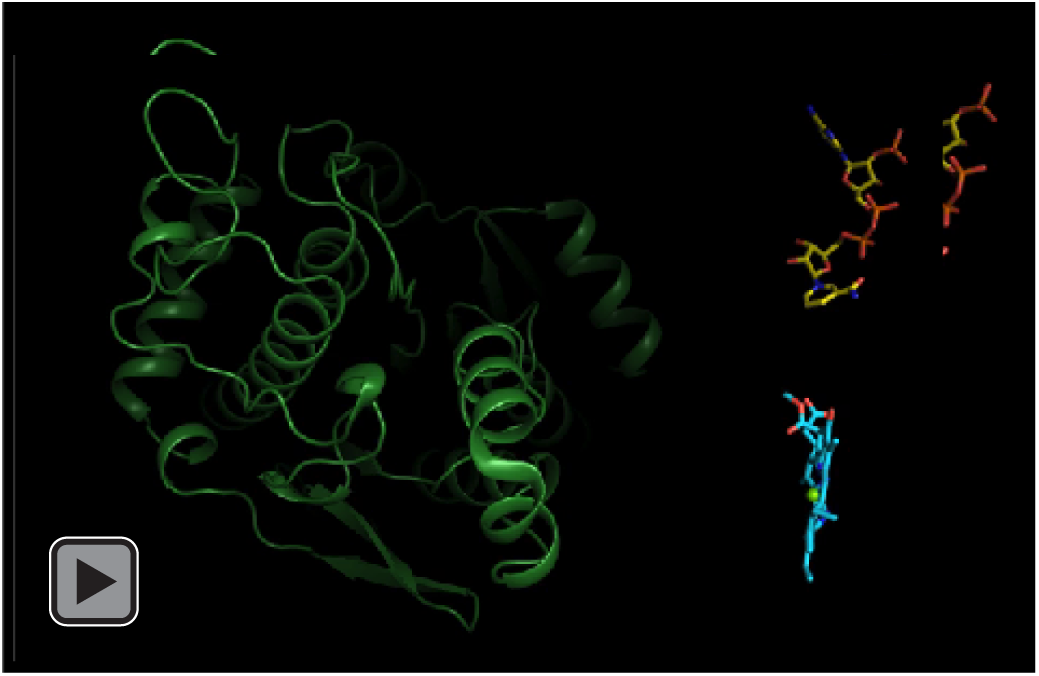
Conformational changes upon Pchlide binding.

### Pchlide induces AtPORB conformational changes that promote insertion into the membrane

To understand the effect of Pchlide and MGDG binding on LPOR, we compared our structure to the Pchlide-free, NADPH-bound cyanobacterial LPORs^6,14^ (Fig. 3d, Fig. S6, Video S 2). The overall root mean squared deviation of Cα atoms ranges from 0.89-0.95 Å. Compared to the cyanobacterial structures lacking Pchlide, however, the Pchlide loop drastically rearranges. In particular, the loop in the *Synechocystis* structure occludes Pchlide binding. In addition, in the *T. elongatus* structure, helix α10 is in a strikingly different conformation. The arrangement of the Pchlide loop and helix α10 in our structure results in the formation of a hydrophobic surface for membrane penetration. Thus, Pchlide binding may induce conformational changes that favor membrane insertion, either followed by or in concert with MGDG binding. LPOR recruitment to the bilayer would increase its local concentration and promote filament polymerization.

### LPOR likely developed unique catalytic residues

The stereo-specific reduction of the C17=C18 double bond of Pchlide by LPOR has been investigated extensively. Hydride transfer from the *proS* site of NADPH^20^ was proposed to reduce C17 first in response to photon absorption by Pchlide, followed by proton transfer from AtPORB Y276 to C18^2^. The involvement of residue Y276 was inferred from homology to other related short chain reductases, where YxxxK is a conserved catalytic motif^21^. Our 3D reconstruction reveals for the first time the architecture of the catalytic site of LPOR and challenges this proposed mechanism (Fig. 3e). Surprisingly, the distance between the NADPH hydride and C17 of Pchlide is ∼4.6 Å, and the attack angle is unfavorable for catalysis. This contrasts with the measured rate of the hydride transfer, which is significantly faster than any other known reported hydride transfer reaction^22^. Such an unexpected orientation of NADPH relative to Pchlide C17 may act to prevent catalysis in darkness, allowing for filament formation and an optimized reaction upon illumination^22^.

Intriguingly, the proposed H^+^ donor, Y276, is far from Pchlide C18 (∼7.3 Å). This distance strongly suggests that it does not play a catalytic role, but is primarily involved in NADPH binding (Fig. 3e). Importantly, homologous mutations of Y276 in cyanobacterial LPOR (Y193F, Y193A, and Y193S) significantly decreased, but did not abolish activity^18^. This partial loss-of-function indicates that either a further structural rearrangement occurs (which is unlikely given the protonation rate is ∼0.2-0.4 10^6^ s^-1^ for plant LPORs^23^), water is involved in the reaction (not resolvable at our resolution), or that there is an alternative donor. We propose that C309 is a possible candidate, given it is ∼4.4 Å away from the C18 and present with the proper dihedral angle for proton transfer (Fig. 3e). Moreover, this residue is highly conserved across kingdoms (Fig. S4b) and a loss-of-function mutation at this position strongly decreases enzymatic activity^24^.

After light-induced catalysis, we speculate that the rearrangement of helix α10 may play a role in the disassembly of the filament. Upon reduction of the C17=C18 bond, the propionic group at C17 gets pushed out of the plane of the ring, allowing the interaction between its carboxyl group and the amide moiety of NAPDH (Fig. 4a). Concurrently, the carboxyl group becomes surrounded by three hydrophobic residues (F316, F323 and F327) that lay on or just upstream of helix α10. We hypothesize that this unfavorable environment promotes rearrangement of the helix outward, similar to the conformation seen in the Pchlide-free state (PDB:6L1G, Fig 4a). This rearrangement would disrupt both the pigment binding and lipid bilayer interactions observed in our structure. Consequently, LPOR could dissociate from the membrane, leaving the product, Chlide, embedded within the outer leaflet for the subsequent and final reaction in Chl biosynthesis by the transmembrane Chl synthase (Fig. 1a, 4b).

**Fig 4.**
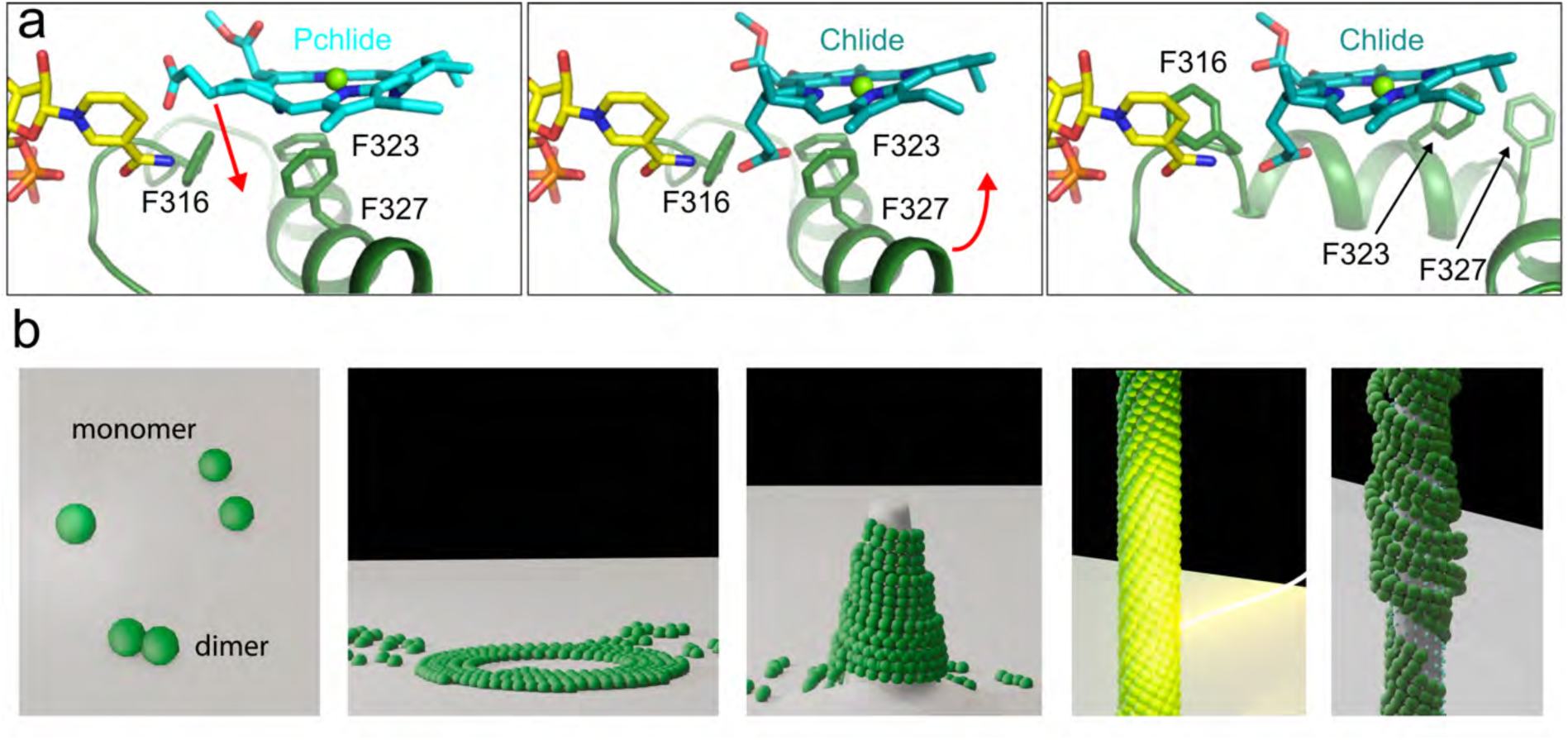
Reduction of Pchlide to Chlide may introduce conformational changes that lead to the filament disassembly. **a**, Conformational mismatch of Chlide in the pigment binding pocket. Left, Pchlide in our reconstruction. The arrow indicates the movement of the propionic group upon stereo-specific reduction of the C17=C18 bond. Middle, Model of the Chlide in our reconstruction. Steric hindrance between the polar carboxyl group of Pchlide and hydrophobic residues of helix α10 may trigger the rearrangement of the helix along the arrow. Right, Model of helix α10 displacement for product release based on PDB:6L1G. **b**, Simplified cartoon model of LPOR functioning in the plant cell.

### Plant LPOR is a dual-purpose enzyme that exploits light-regulated dynamics to form membranes in chloroplasts

Our 3D reconstruction shows how plant LPOR binds to Pchlide, NADPH, and lipids to form filaments that remodel membranes. This self-assembly facilitates efficient energy transfer for the reduction of Pchlide, while simultaneously targeting the product to the bilayer (Video S 3). By coupling our structural analysis with detailed sequence comparisons of hundreds of plant and cyanobacterial LPOR sequences, we identify evolutionary differences in higher-order oligomerization interfaces (Fig. S4). The AtPORB residues involved in oligomerization, as well as the hydrophobic nature of the Pchlide loop and helix α10, are highly conserved across plants, implying that they have evolved to form lipid-shaping filaments in a light-dependent manner. We recently showed that plant LPORs, in contrast to cyanobacterial counterparts, have a unique PG-driven NADPH binding mechanism and MGDG-dependent spectral properties^15^. Altogether, this raises new questions about the physiological role of such lipid-specific adaptations.

The filaments are astonishingly similar to elements of the PLB, as both are composed primarily of the same lipids and have the same spectral properties and similar cylindrical dimensions. However, intact PLBs are a 3D cubic lattice of interconnected tubules, while our in vitro assemblies are long cylinders only. Strikingly, however, high salt, lower pH and freeze-thaw treatment dissolve PLBs into similar tubules ^25^. Based on these observations, we believe PLBs are composed of LPOR filaments that, likely with other factors, form the 3D organized lattice.

We reveal that plant LPOR forms helical filaments with lipids from the inner membrane of both chloroplasts and PLBs, as long as there is Pchlide, NADPH, and darkness (Fig. S1). This naturally occurs during etiolation (i.e. when germinating underground or in leaf buds), but also at night in developed leaves. In etiolated tissue, PLBs, the protein constituent of which is primarily LPOR, provide efficient Chl biosynthesis as well as the lipids for newly forming photosynthetic membranes^4,9^. Given that AtPORB is expressed in both mature and etiolated leaves, we hypothesize that tubule formation by LPOR may also take place in mature leaves during the night, and could therefore provide a reservoir of lipids for new thylakoids^26^. A mechanism for lipid trafficking between the chloroplast inner membrane and the thylakoids was postulated but has not been discovered^27,28^. In support of our conjecture, the *A. thaliana porB/porC* double mutant lacks stacked thylakoids^29^. This phenotype implicates a role for LPORs in thylakoid formation. LPOR oligomerization may, therefore, play a role beyond light-dependent chlorophyll precursor biosynthesis by providing a mechanism for sunlight to regulate lipid availability and organization in chloroplasts.

**Video S3.**
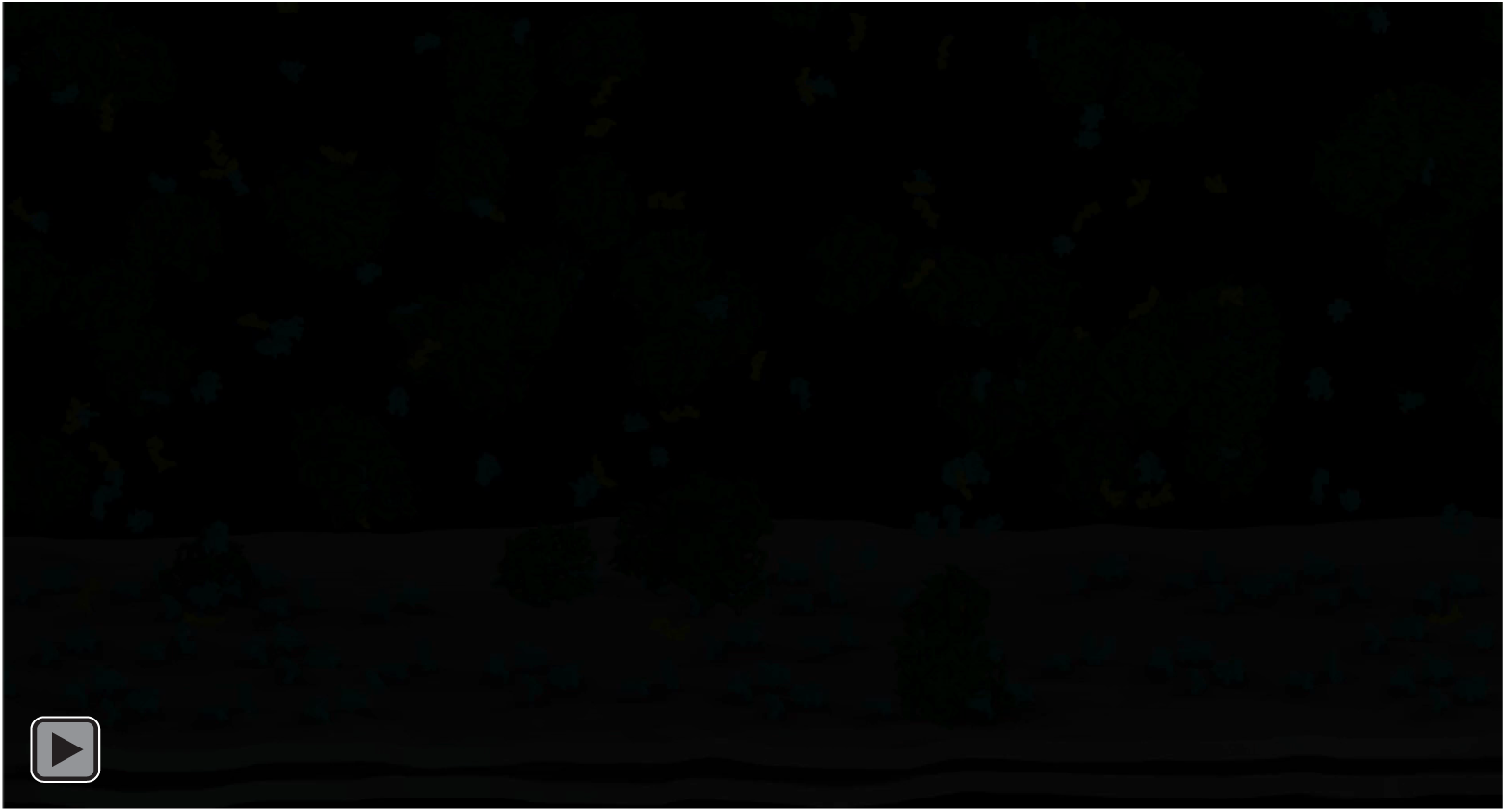
Mechanism of substrate binding, polymerization, and membrane remodeling by plant LPOR.

## Acknowledgements

We thank: the Frost lab for helpful discussions and critical comments; P. Thomas, L. E. Miller-Vedam, and M. Sun with computational assistance; N. Poweleit for discussions on helical processing; T. Borowski for generating Pchlide ligand restraints; M. Braunfeld, D. Bulkley, M. Harrington, A. Myasnikov, and Z. Yu of the UCSF Center for Advanced CryoEM for microscopy support; and J. Baker-LePain and the QB3 shared cluster (NIH grant 1S10OD021596-01) for computational support. Structural biology applications used in this project were compiled and configured by SBGrid. The Titan X Pascal used for this research was donated by the NVIDIA Corporation. This study was supported by a Bekker scholarship granted by the Polish National Agency for Academic Exchange (NAWA) PPN/BEK/2018/1/00105 and START scholarship granted by the Foundation for Polish Science (FNP) 024.2018 (to M.G) and NIH 1DP2GM110772-01 (to A.F). A.F. is further supported by a Faculty Scholar grant from the HHMI and is a Chan Zuckerberg Biohub investigator.

## Author contributions

M.G. and A.F. conceived of the project. M.G., A.F., and H.C.N. designed experiments and analyzed data. M.G. optimized and produced the samples, performed spectroscopy measurements, and sequence alignment analysis. H.C.N. assisted in sample optimization and performed cryoEM experiments and analysis. J.K. and M.G. purified Pchlide. A.A.M. prepared the animation video and assisted with cryoEM data collection. H.C.N., A.F., and M.G., prepared the manuscript with input from all authors.

## Methods

### Protein expression and purification

AtPORA, AtPORB, AtPORC, and *Synechocystis* LPOR were expressed in *E. coli* BL21(DE3)pRIL strain with N-terminal HisTag as previously described^30,31^.

### Protochlorophyllide purification

Protochlorophyllide was isolated from etiolated wheat seedlings as previously described^31^.

### Low-temperature fluorescence spectra measurements

Low-temperature (at 77 K) fluorescence measurements were performed and analyzed as previously described^31^.

### Filament assembly

MGDG, PG, and DGDG (50 mol%, 15 mol%, 35 mol%; Lipid Products) in chloroform:methanol (2:1 v/v) solution were mixed in a glass vial and the solvent was evaporated under a stream of nitrogen. The lipid film was then dried under vacuum for 30 minutes, hydrated and vortexed in arginine-rich buffer (0.6 M arginine, 50 mM phosphate buffer, 50 mM NaCl, 7mM 2-mercaptoethanol, pH 7.1). The final total lipid concentration was 2 mM.

To set up the filament assembly, 30-60 μM of the freshly prepared lipid films were resuspended in low-arginine buffer (0.18 M arginine, 50 mM phosphate buffer, 50 mM NaCl, 7mM 2-mercaptoethanol, pH 7.1). Then, 800 μM NADPH and 8-18 μM LPOR were added, so the total volume of the sample was 100 μl. All the following procedures were conducted in darkness or under dim green light that was previously shown not to trigger the reaction^32^. To initiate filament assembly, 10 μM Pchide was added and the reaction mixture was immediately vortexed for 2 seconds. The sample was incubated at room temperature in darkness for 5 minutes, and then centrifuged for 5 minutes (16,000 g). Next, the supernatant was carefully removed, and the remaining intensely green pellet was resuspended in 10 μl of arginine-rich buffer. The filament sample was protected from light until stained with uranyl acetate or frozen in liquid ethane.

### EM sample preparation and data acquisition

For negative stain EM imaging, 7 μl of the filament sample were first applied to a glow-discharged grid covered with carbon film for 1 minute and then removed by blotting with filter paper. The grid was then washed once in water and then stained with 2% (w/v) uranyl acetate. Grids were imaged with a Tecnai T12 microscope (FEI) operating at 120 kV equipped with a Gatan Ultrascan CCD camera (Gatan). For cryoEM imaging, 3.5 µL of the assembled filament were applied to glow-discharged R1.2/1.3 Quantifoil 200 Au mesh grids (Quantifoil) in a Mark IV Vitrobot (FEI). Grids were blotted with Whatman #1 filter paper (Whatman) for 4-8 seconds with a 0 mm offset at 19 °C and 100 % humidity before plunging into liquid ethane. Grids were stored under liquid nitrogen until samples were imaged for structural determination. The dataset was collected on a 300 kV Titan Krios operating with C2 aperture of 70 μm and an energy filter with a 20 eV slit using a K3 direct detector operated in super-resolution mode and binned by a factor of 2 for subsequent processing (resulting Å/px of 0.835). Data collection parameters are summarized in Table S1.

### EM Image Analysis and 3D Reconstructions

Structural biology applications used in this project were compiled and configured by SBGrid^33^. All dose-fractionated image stacks were corrected for motion artifacts, 2x binned in the Fourier domain, and dose-weighted using MotionCor2^34^. GCTF-v1.06^35^ was used for contrast transfer function (CTF) estimation. Filaments were autopicked with crYOLO^36,37^, using a manually trained model on 50 micrographs for filament mode and a 512 px box. Subsequent steps were performed in RELION3^38^ unless otherwise stated. Segments were extracted with ∼90% overlap between boxes and binned 4x to a 128px box. The ‘ignore CTFs until first peak’ flag for CTF estimation was used throughout. Multiple rounds of 2D classification were performed to remove poor particles and to yield particles with a largely uniform diameter and well-defined features. The segments were then binned 2x (256 px box) and a hollow, smooth cylinder was used as the initial model for the first round of 3D auto-refine without applying helical symmetry. This led to an initial reconstruction with some definable subunit positions. These particles then were subject to 3D classification without applying helical symmetry to identify and remove heterogeneous diameters. The classes with well-defined secondary structures were then subject to a round of 3D auto refinement still without applying helical symmetry. This reconstruction yielded better-defined subunits and was used to search for helical parameters. 3D classification was then performed with refinement of the helical parameters and a central Z length of 40% of the particle box. The best particles then went through multiple rounds of 3D classification without alignment and Tau^2^ values varying from 2-10 with a protein-membrane mask. Selected segments were then unbinned to a 384 px box (∼1.3x, resulting Å/px of 1.113333) and subject to another 3D auto-refine reconstruction. Iterative cycles of per-particle beam tilt/trefoil and magnification anisotropy CTF refinement were performed with 3D auto-refinement. Unbinning to the full 512 px box did not improve the reconstruction. Bayesian polishing was performed but did not improve the reconstruction likely due to the few number of particles per micrograph. Given the relatively large asymmetric unit, a final 3D auto-refine run was performed using local symmetry within RELION encompassing 54 promoters, the number of complete protomers within 40% of the central Z length. Postprocessing and half map Fourier shell correlations (FSCs) were performed on the half maps without any externally imposed local or helical symmetry. The map used for atomic modeling had local symmetry and then helical symmetry applied using a central Z percentage of 40%. The refined helical parameters are a twist and rise of 50.34° and 43.1 Å, respectively. The reconstruction has C2 helical point group symmetry. Refinement parameters are listed in Table S1.

### Atomic Modeling and Validation

A homology model of PORB was generated using the I-ITASSER^39-41^ web server. This monomer was initially docked into the EM map with UCSF Chimera^42^. The density surrounding the monomer was extracted using phenix.map_box^43^ for use for manual refinement *Coot*^44^ and subsequentphenix.real_space_refine^43^ using global minimization, morphing, secondary structure restraints, and local grid search. The refined protomer were then used to generate nine surrounding protomers from the local and helical symmetry applied post-processed map using Chimera. Noncrystallographic symmetry (NCS) constraints were then used throughout refinement in phenix.real_space_refine. Iterative cycles of manually rebuilding in *Coot* and phenix.real_space_refine with previous strategies and additionally B-factor refinement were performed. The final model was then built containing 40 protomers using phenix.apply_ncs.

All final model statistics were tabulated using Molprobity^45^ and summarized in Table S1. Half map and map versus model FSC plots were computed in PHENIX^46^. All structural figures were generated with UCSF ChimeraX^47^ and PyMOL (www.pymol.org). The final animation was generated in Blender (www.blender.org).

### Phylogenetic analysis and sequence similarity analysis

The LPOR sequences were analyzed as previously described^31^.

### Report Summary statement

Further information on experimental design is available in the Nature Research Reporting Summary linked to this article.

### Data Availability

The atomic model and cryoEM maps were deposited to the PDB (7JK9) and EMDB (EMD-22364).

